# Time in sympatry correlates with the strength of reproductive isolation in hybridising *Typha*

**DOI:** 10.1101/2025.09.12.675895

**Authors:** Alberto Aleman, Joanna Freeland, Marcel Dorken, Aaron Shafer

**Author notes:** **Corresponding authors:** Aaron Shafer, Alberto Aleman.

## Abstract

In North America, hybridisation between *Typha angustifolia* and *T. latifolia* results in the highly impactful *T.* × *glauca*. In Europe, where its parental taxa are also sympatric, *T*. × *glauca* is scarce, suggesting stronger reproductive isolation on this continent. Using genomic data from the two species in North America and Europe, we reconstructed their demographic histories and characterised barrier loci between them. Demographic modelling suggests that the initial contact between the two species in Europe occurred ∼800,000 years ago, indicating sympatry in that region since the Middle Pleistocene. In North America, their contact likely happened within the last centuries and was potentially driven by the human-mediated dispersal of *T. angustifolia*. We identified 47 candidate barrier loci between species in Europe, 6 of which are associated with reproductive functions. No barrier loci were found in North America. Our results suggest that prolonged sympatry can promote the evolution of reproductive barriers, whereas prolonged allopatry can reduce the likelihood of their development. Future studies could help determine whether time in sympatry is a predictor of the strength of reproductive isolation between hybridising taxa. Preventing invasions by hybrid taxa will require limiting the human-mediated dispersal of *Typha* (and other allopatric species) lacking reproductive isolation.

## Introduction

Hybridisation is an important evolutionary mechanism (Abbott et al. 2013); it can cause the emergence of new species if reproductive barriers develop between parental and hybrid lineages, or lead to genetic swamping and the extinction of parental taxa (Runemark et al., 2019). More commonly, hybridisation results in the formation of hybrid zones—geographic areas where parental and hybrid individuals of one or more generations coexist (Hewitt, 1988). Hybrid zones provide an opportunity to characterise the genetic architecture of interspecific gene flow and reproductive isolation (Taylor & Larson, 2019)

The formation and persistence of hybrid zones are influenced by species boundaries (Larson et al., 2014). Different types of reproductive barriers maintain these boundaries (Harrison & Larson, 2014), with “barrier loci” referring to genetic regions that underlie the traits controlling reproductive isolation between species (Barton & Bengtsson, 1986). Barrier loci can act prezygotically by impeding interbreeding events (Elmer, 2019), or postzygotically by making hybrids sterile, inviable, or unfit (Stankowski & Ravinet, 2021). Prezygotic barrier loci are often under divergent selection (Wu, 2001). For instance, in *Heliconius* butterflies, the optix and WntA wing patterning loci govern mate recognition and limit interspecific courtship (Merrill et al., 2019).

Postzygotic barrier loci frequently arise from intrinsic epistatic incompatibilities, as those explained by the Bateson–Dobzhansky–Muller (BDM) model, in which mismatched allele combinations from different loci decrease hybrid fitness (Bateson, 2009; Dobzhansky, 1936; Muller, 1942); in *Icterus* orioles, a large inversion on the Z chromosome accumulates alleles that cause hybrid breakdown (Walsh et al., 2023). Understanding the role of barrier loci to reproductive isolation is key to explaining how species boundaries evolve.

Different approaches have been used to investigate the loci that affect reproductive isolation between taxa (e.g., Moyle and Payseur 2009; Schumer et al. 2014; Haenel et al. 2021). In recent years, methods for identifying barrier loci from genome-wide sequencing data have been developed (Burban et al., 2024; Laetsch et al., 2023).

These methods use coalescent-based simulations and explicitly incorporate demographic inferences (i.e., divergence times, population sizes, migration rates) (Fraïsse et al., 2021). Demographic-informed approaches to identify barrier loci overcome two limitations of methods based on genomic scans: (i) that the genomic landscapes of diversity and differentiation (e.g., F_ST_, d_XY_, π) can emerge from evolutionary forces not directly related to reproductive isolation (e.g., historical bottlenecks, genetic drift, local adaptations), and (ii) that there is no universal summary statistic for barrier loci, as the emergence of these barriers depends on multiple factors, reflected on multiple summary statistics. Identifying barrier loci through demography-aware methods can provide insights into how reproductive isolation evolves throughout species’ histories.

Across the Great Lakes, Prairie Pothole, and Midwestern regions of North America, the hybrid cattail *Typha × glauca* is a highly impactful invader (reviewed in Bansal et al. 2019). Hybrid cattails outcompete their parental species—*T. angustifolia* and *T. latifolia*—(Freeland et al., 2013, 2024; Geddes et al., 2021; Pieper et al., 2020; Zapfe & Freeland, 2015), reduce biodiversity and alter freshwater functions (Angeloni et al., 2006; Boers et al., 2007; Farrer & Goldberg, 2014; Tuchman et al., 2009), and are expanding across their already broad range (Joyee et al., 2024). Contrastingly, *T. × glauca* is scarce in Europe and has not been reported in Asia, where its parental species also co-occur (Ciotir et al., 2017; Nowińska et al., 2014; Volkova & Bobrov, 2022; Zhou et al., 2016). A key difference in the history of both parental species in North America and Europe is a longer time inhabiting Europe, where they might have been sympatric since the emergence of the younger, *T. latifolia*, ∼5.7 Ma (Zhou et al., 2018), or once it recovered from a severe demographic decline and subsequently expanded ∼800,000 years ago (Aleman et al., 2025). *Typha latifolia* arrived in North America via the Beringian Land Bridge from Eastern Eurasia—the centre of origin of *Typha*—between the Late Miocene and Early Pliocene, i.e., from 5.7 to 3.5 Ma (Zhou et al., 2018), while *T. angustifolia* likely arrived centuries ago, potentially due to human dispersal (Ciotir et al., 2013).

Although we cannot rule out the possibility that the two species might have experienced contact in East Eurasia before *T. latifolia* arrived in North America, the available information indicates that *T. latifolia* has been isolated from *T. angustifolia* since it entered North America and until *T. angustifolia* was introduced (Ciotir et al., 2013; Ciotir & Freeland, 2016).

The broad distribution of the *T*. × *glauca* hybrid swarm in North America suggests that reproductive barriers between *T. latifolia* and *T. angustifolia* are weak on this continent. In contrast, the scarcity of *T*. × *glauca* in Europe suggests that there are strong reproductive barriers between its two parental species in this region. Aleman et al. (2025) identified introgressive hybridisation from *T. latifolia* to *T. angustifolia* in North America and Europe, which, together with the scarcity of *T*. × *glauca* in Europe, suggests that the two species hybridised in the past in Europe and reproductive barriers emerged later.

We hypothesised that (i) in North America, the time in isolation that *T. latifolia* experienced before *T. angustifolia* entered the continent reduced the possibility for natural selection to drive the evolution of reproductive isolation between the two species, whereas (ii) in Europe, their longer coexistence and historical hybridisation led to the development of reproductive barriers. Therefore, we expected to identify more barrier loci between the two species in Europe than in North America. Characterising these loci could help inform why *T*. × *glauca* is a highly impactful invader in some parts of North America but not in Europe. Understanding what limits hybridisation between the two species in Europe should receive special consideration, since *Typha* is frequently transported between continents via garden centres (Ciotir & Freeland, 2016). If barriers to hybridisation are genetic—rather than ecological—the human-mediated movement of *Typha* could lead to the contact of species lacking reproductive isolation, facilitating future invasions by novel hybrids. From a theoretical standpoint, the time in sympatry for *T. angustifolia* and *T. latifolia* in North America and Europe is uniquely positioned to test whether barriers to hybridisation could be stronger between taxa that have experienced prolonged periods of contact than those whose time in sympatry is very recent.

## Materials and Methods

### Genomic data processing

Raw sequencing data from *T. angustifolia* (36 from North America, 16 from Europe) and *T. latifolia* (47 from North America, 23 from Europe) were obtained from Aleman et al. (2025) and supplemented with data from additional samples (Figure 1; Supplementary Table S1), generated following Aleman et al., (2024). The quality of the demultiplexed raw sequences was evaluated using FastQC 0.11.9 (Andrews, 2017) and MultiQC 1.14 (Ewels et al., 2016). Read pairing and adapter trimming were performed using Trimmomatic 0.39 (Bolger et al., 2014), removing reads shorter than 100 bp after trimming. Cleaned reads were mapped to the *T. latifolia* nuclear (GenBank accession JAIOKV000000000.2 (Widanagama et al., 2022)) and chloroplast (GenBank accession NC_013823 (Guisinger et al., 2010)) genomes using BWA 0.7.17 (Li & Durbin, 2009). Mapping statistics were generated using SAMtools 1.15.1 (Li et al., 2009). Genotyping was conducted using ANGSD 0.93 (Korneliussen et al., 2014). SNPs were called for all samples, requiring minimum mapping and sequencing scores of 20, as well as a minimum *p*-value of 1^e−6^. Sites mapped to the plastome were removed using VCFtools 0.1.16 (Danecek et al., 2011). Two SNP datasets were generated based on two missing data thresholds across all samples (i.e., without considering taxonomy or geographic location): 20% for the genetic structure analysis, to ensure high-confidence genotypes for accurate sample classification (Yi & Latch, 2022), and 50% for the demographic history and barrier locus characterisation, to maximize SNP retention, since coalescent-based methods explicitly model genotype uncertainty and require larger datasets for robust parameter estimation (Fraïsse et al., 2021). To confirm the samples’ species membership, their genetic relationships were evaluated by a neighbour-joining (NJ) tree analysis, conducted using Plink 1.9 (Purcell et al., 2007).

**Figure 1.**
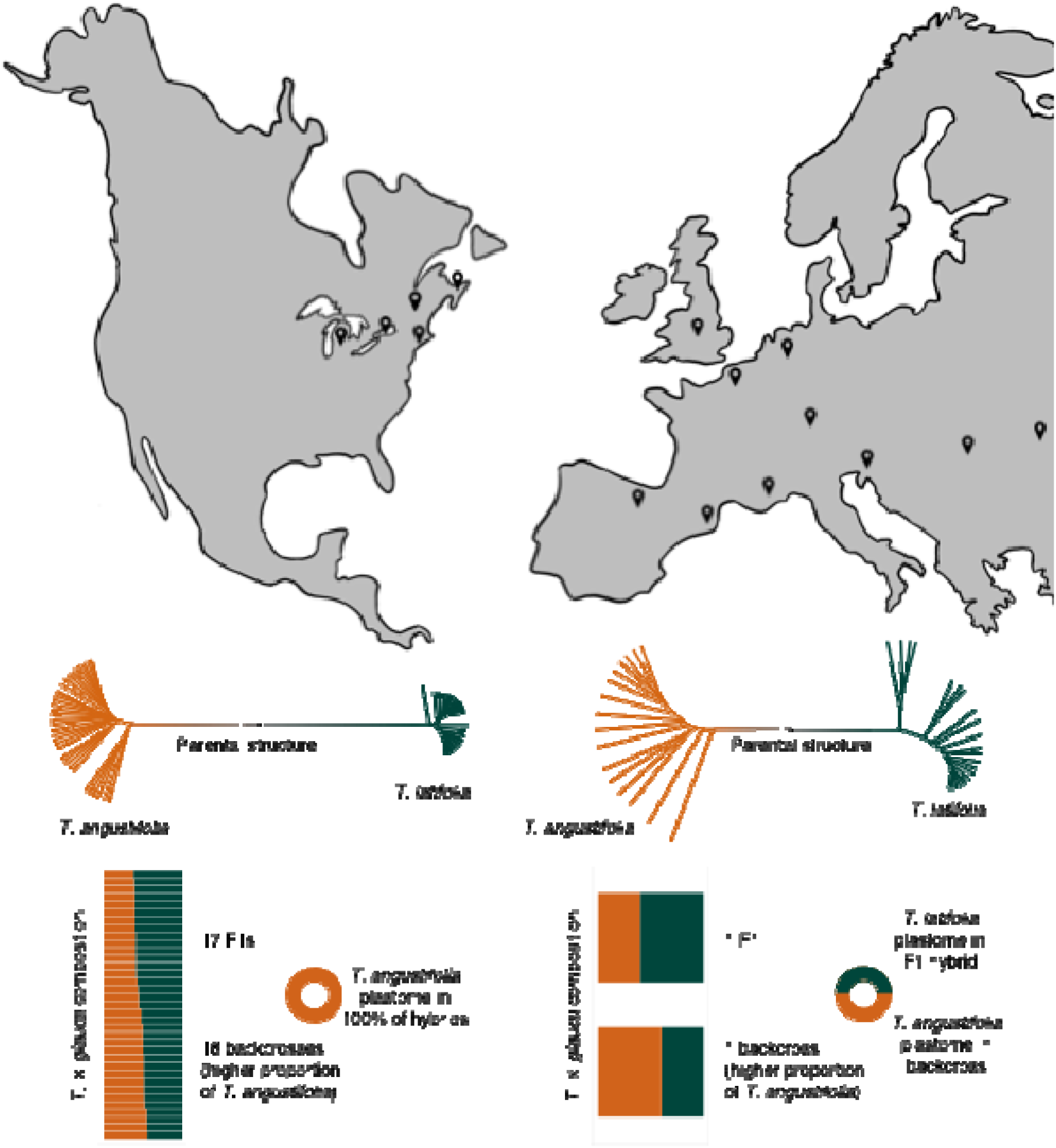
Top: Approximate sampling sites (black marks) in this study; sample sizes are not represented graphically. Middle: NJ trees for *Typha angustifolia* and *T. latifolia* in this study; branches represent samples, and colours indicate species, as labelled. Bottom: Genetic composition of *T. × glauca* samples across the hybrid zone.

### Genetic composition of the T. × glauca hybrid swarm

To assess the genetic composition of hybrids across the *T. × glauca* swarm, sequencing data were generated from 38 hybrid samples following Aleman et al. (2024). Thirty-six of these samples were from North America, whereas only two were found in Europe, despite an extensive collecting trip that aimed to maximise the number of sampled hybrids (Ciotir et al., 2017). In North America, the 36 hybrids were collected during two previous studies (Pieper et al., 2020; Tisshaw et al., 2020) (Figure 1; Supplementary Table S1). Raw sequences were quality-inspected, cleaned, and mapped to the *T. latifolia* reference genome following the *Genomic data processing* steps above.

SNPs were called for the 42,167 sites (see *Results*) used in the genetic structure analysis for *T. angustifolia* and *T. latifolia*. The nuclear composition of hybrids was assessed using ADMIXTURE 1.3.0 (Alexander & Lange, 2011), which tested *K* clusters from 1 to 10 for all samples in this study. The optimal *K* number was chosen via a cross-validation procedure. To assess whether hybrids had the plastome of *T. angustifolia* and *T. latifolia*, the plastomes of all samples in this study were assembled using the reference-based approach in Aleman et al. (2024), and SplitsTree4 (Huson & Bryant, 2006) was used to reconstruct a phylogenetic network for all samples in this study.

### Demographic history and characterisation of barrier loci

To reconstruct the demographic history of *T. angustifolia* and *T. latifolia* in North America and Europe, and to identify barrier loci between the two species, RIDGE (Burban et al., 2024) was run between species within and between continents (e.g., North American *T. angustifolia* with European *T. latifolia*); i.e., four independent analyses were conducted (Table 1). No hybrids were included in these analyses. Before the demographic inferences, several summary statistics were computed in 5 kb windows, including the genetic differentiation between species through F_ST_ (Hudson et al., 1992), d_XY_ (Nei & Miller, 1990), d_a_ (Nei & Li, 1979), and their joint Site Frequency Spectrum jSFS (Wakeley & Hey, 1997), and the diversity within each species through π (Nei & Li, 1979), Watterson’s θ (Watterson, 1975), and Tajima’s D (Tajima, 1989). Four demographic scenarios of divergence between species were simulated using an Approximate Bayesian Computation framework: strict isolation, ancestral migration, secondary contact, and isolation–migration. These scenarios included both homogeneous and heterogeneous population sizes, as well as interspecific gene flow rates over time (Figure 2). The prior distributions of species’ divergence times and effective population sizes were determined based on the summary statistics observations (Supplementary Table S2). To avoid having more than two orders of magnitude between the prior bounds of the time at which the two species diverged from a common ancestor, the minimum divergence time was adjusted to two-thirds of the estimated maximum divergence time. However, the priors for the minimum and maximum population sizes were left unadjusted. During each of the four independent analyses, 7.5 M sites were computed over 560 simulations.

**Figure 2.**
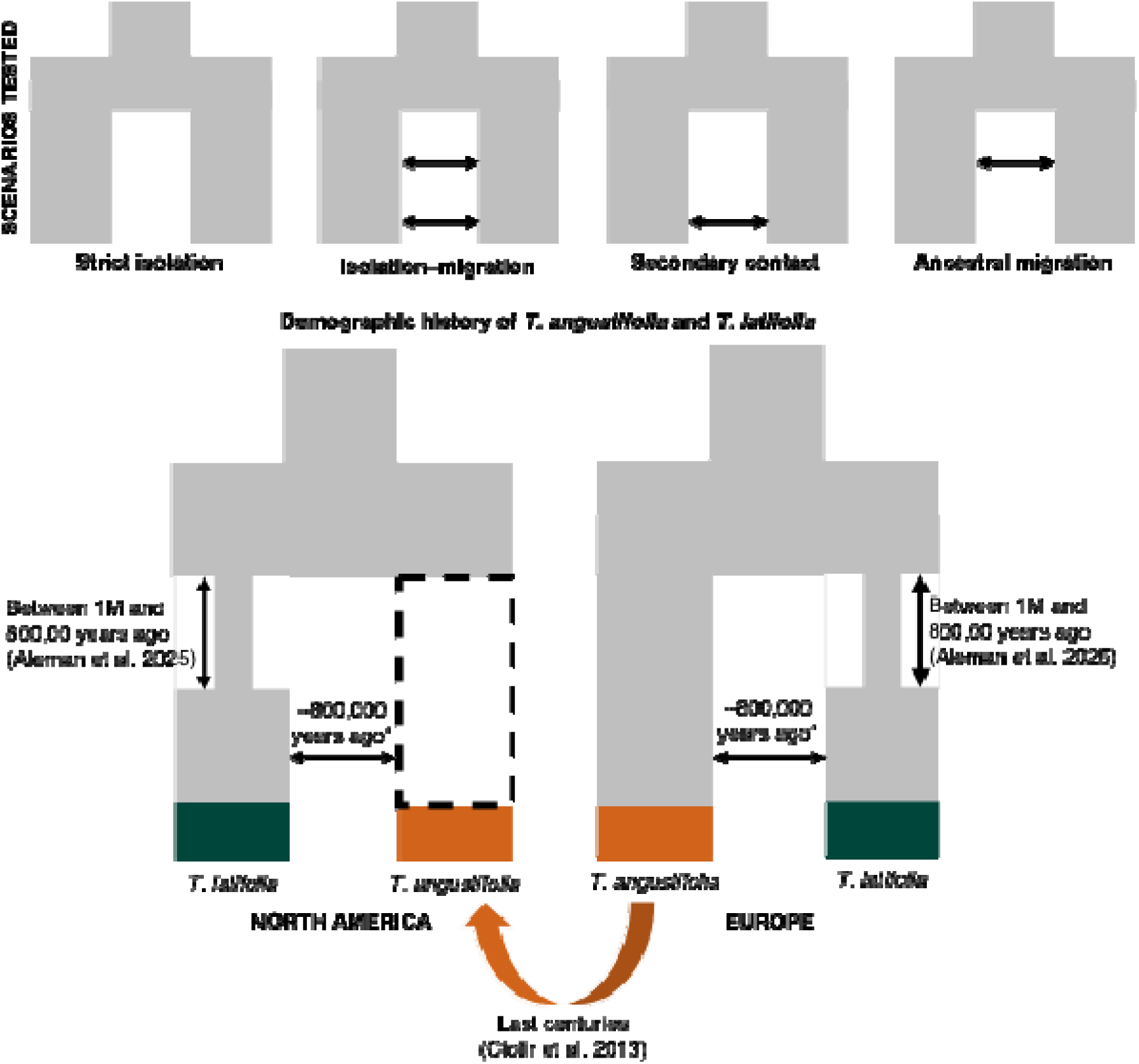
Demographic history of *T. angustifolia* and *T. latifolia* in North America and Europe proposed in this study. *Parameter estimated by RIDGE in this study.

**Table 1.**
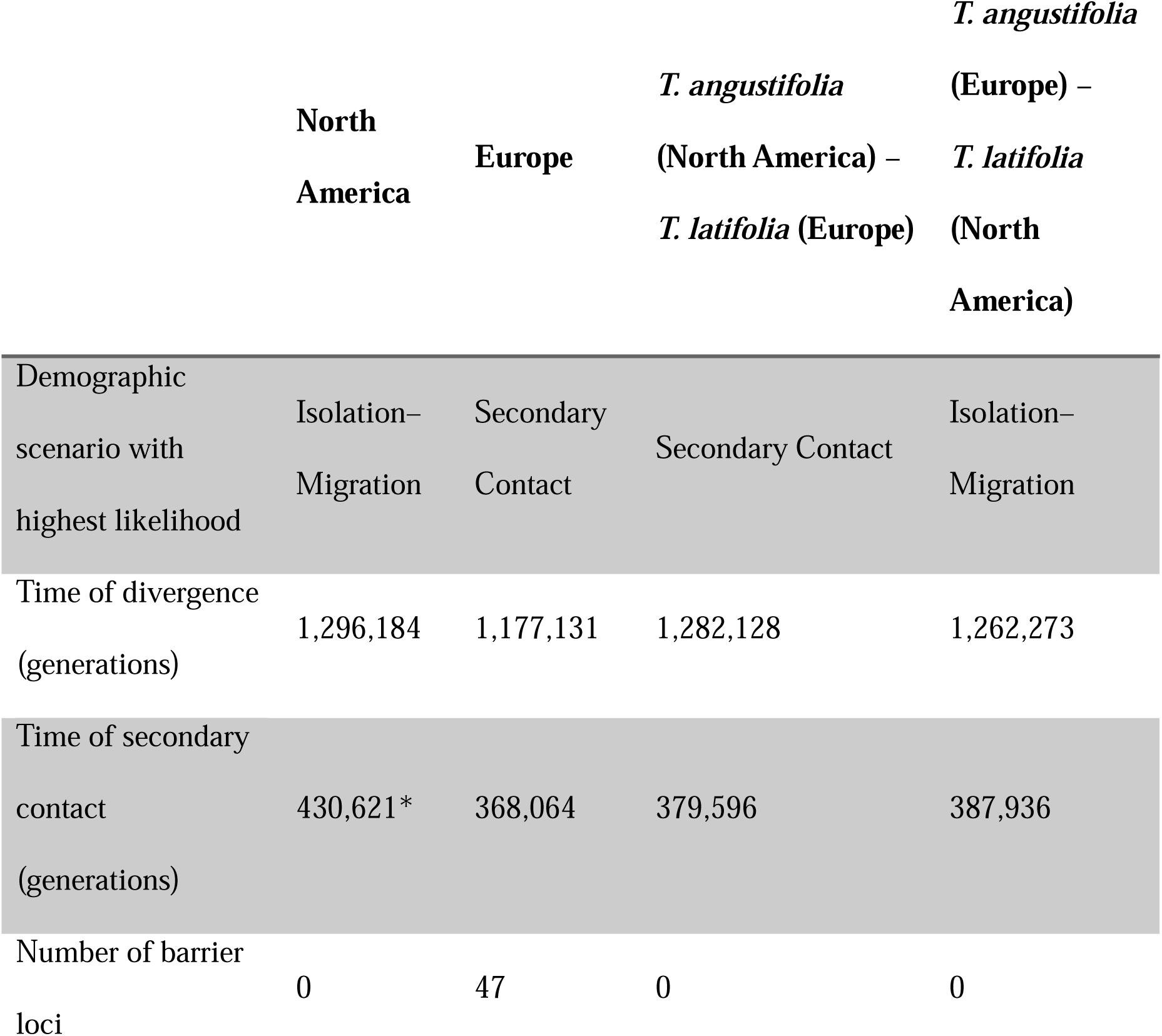
Demographic history of *T. angustifolia* and *T. latifolia* in North America and Europe. Parameters estimated by RIDGE. *Ciotir et al. (2013) showed that *T. angustifolia* potentially arrived in North America within the last centuries, which is the accepted time of secondary contact between the two species.

Simulation results were used to build a demographic hypermodel, where each parameter was obtained from the weighted likelihood of the four scenarios tested. The hypermodel was used to simulate 1.5 M barrier and non-barrier loci, i.e., with *m* = 0 and *m* > 0, respectively, where *m* is the effective gene flow rate between species. The probability that a simulated locus was accurately identified as a barrier or non-barrier locus was estimated to obtain a false-positive and false-negative correction (*Q*). The simulation results were used to calculate the likelihood that a real-life locus was a barrier to hybridisation. The identification of a locus (i.e., a 5 kb window) acting as a barrier to hybridisation was determined by a Bayes Factor of 10, given the correction *Q* (known as “BF_approxQ”); this meant that a barrier locus was at least 10 times more probable to be a barrier to hybridisation than a non-barrier locus. Barrier loci identified in each run were compared, and summary statistics for barrier and non-barrier loci were evaluated.

BEDTools Intersect (Quinlan & Hall, 2010) was used to identify the genes associated with barrier loci from the *T. latifolia* annotation (Widanagama et al., 2022). The ratio of nonsynonymous to synonymous substitution rates (dN/dS) for all barrier loci was calculated among the four groups of taxa (i.e., the two species from the two continents) using MEGA 12 (Kumar et al., 2024). Barrier loci that could not be associated with genes from the *T. latifolia* annotation were screened using BLAST+ to search for homologous genes (Camacho et al., 2009).

## Results

Using 42,167 SNPs, the NJ tree confirmed that in each continent, parental samples clustered into two groups (Figure 1). The ADMIXTURE analysis, including parental and hybrid samples, established the most likely number of genetic clusters as two (*K* = 2). In North America, 19 hybrids were F1s and 17 were backcrosses with T. *angustifolia*, with a nuclear proportion of ∼70% from this species; all hybrids (F1s and backcrosses) had the *T. angustifolia* plastome. In Europe, one hybrid was an F1 and had the *T. latifolia* plastome, and the other was a backcross with both a higher nuclear proportion (70%) and the *T. angustifolia* plastome (Figure 1).

The demographic history and barrier locus analyses used 6,735,717 SNPs. In each of the four demographic history reconstructions of *T. angustifolia* and *T. latifolia*, secondary contact and isolation–migration received the highest likelihood weights; these scenarios consistently indicated heterogeneous rates of interspecific gene flow and population sizes over time (Figure 2, Table 1, Supplementary Table S3). The four reconstructions successfully fitted the observed data, with a posterior goodness-of-fit value greater than 0.05. The estimates of divergence and contact times were consistent across all analyses. The mean divergence time between the two species was estimated to be ∼1.25 M generations ago, and the mean time of secondary contact was ∼400,000 generations ago.

A total of 49 barrier loci between *T. angustifolia* and *T. latifolia* were identified, 47 of those in Europe (Figure 3). No barrier loci were identified in North America. Two barrier loci were found between North American *T. angustifolia* and European *T. latifolia* (one of those was also a barrier locus in Europe), and two barrier loci were found between European *T. angustifolia* and North American *T. latifolia* (one of those was also a barrier locus in Europe).

**Figure 3.**
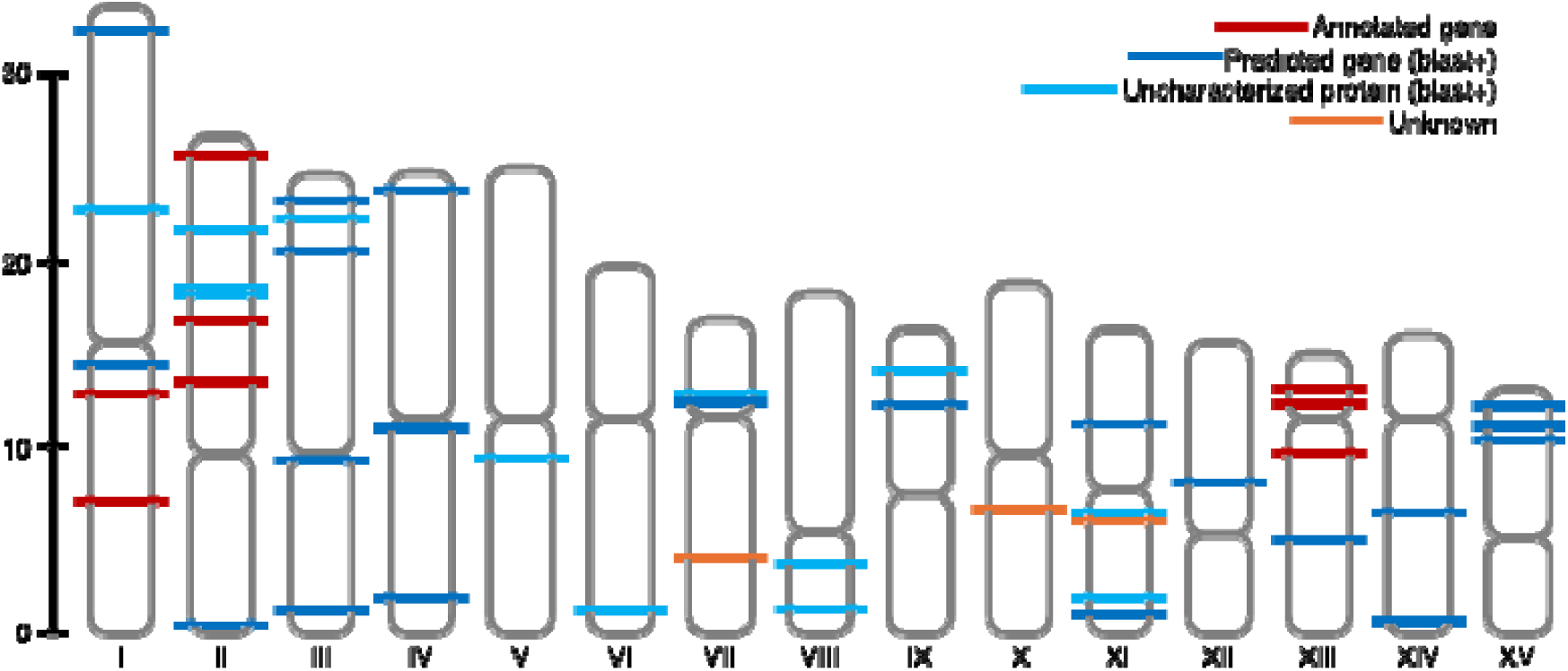
Distribution of loci predicted to be responsible for reproductive isolation between *T. angustifolia* and *T. latifolia* in Europe in this study. Centromeres were predicted using RepeatObserver (Elphinstone et al., 2025) with the default settings.

Of the total 49 barrier loci, 12 were annotated to 10 genes (3 were to the same gene) from the *T. latifolia* reference, and BLAST predicted other 34 to be genes (Supplementary Table S4). In the 47 barrier loci from Europe, European *T. angustifolia* had a mean diversity (π = 0.0028) lower than the rest of its genome (π = 0.0073), and no diversity differences were observed the other three groups of taxa; 10 barrier loci were highly differentiated for F_ST_ between species with values above the 95th percentile, of those, 1 of those was also highly differentiated for d_XY_ and 3 were genes; of the other 7 genes found in barrier loci, 1 was under purifying selection between species in Europe (dN/dS < 1), and 1 was under positive selection (dN/dS > 1) in all groups of taxa except European *T. angustifolia*.

## Discussion

The number and identity of loci associated with species’ reproductive isolation can shed light on how species boundaries evolve along their genomes and through time (Gavrilets, 2003; Ravinet et al., 2017). Additionally, understanding the evolutionary histories of the parental species of *T*. × *glauca* and the mechanisms that promote or limit their interbreeding can help us prevent further invasions by novel *Typha* hybrids. This study provides insights into the demographic histories and barriers to hybridisation between *T. angustifolia* and *T. latifolia*; our results suggest that in Europe, where very few hybrids have been identified, secondary contact between the two parental species occurred ∼400,000 generations ago. *Typha angustifolia* likely entered North America within the last centuries, where it interbreeds with *T. latifolia*, and their hybrids are generally invasive.

Hybridisation between *T. angustifolia* and *T. latifolia* in North America is highly asymmetric, with *T. angustifolia* often being the maternal parent (Ball & Freeland, 2013; Pieper et al., 2017). Consistent with this, the 19 North American F1 hybrids in this study had the *T. angustifolia* plastome. Since hybrid classes go beyond F1, the *T*. × *glauca* hybrid zone is a swarm; F1s can backcross bidirectionally with *T. angustifolia*, asymmetrically with *T. latifolia* as the paternal parent, or reproduce with other F1s (Bhargav et al., 2022; Freeland et al., 2013). Pieper et al. (2017) observed that no seeds are set when *T. latifolia* is pollinated by hybrid cattails. In line with this, we identified 17 backcrosses with a higher nuclear proportion and the plastome of *T. angustifolia*; this observation suggests that *T. angustifolia* is almost always present during backcrossing events. Asymmetric hybridisation can reflect early reproductive isolation, as seen in *Rhododendron delavayi* and *R. cyanocarpum* azaleas (Ma et al., 2016), *Primula vulgaris* primroses and *P. veris* cowslips (Keller et al., 2021), or *Erysimum mediohispanicum* and *E. nevadense* wallflowers (Abdelaziz et al., 2021). However, across the North American Great Lakes, Prairie Pothole, and Midwest, hybrids are generally invasive and outnumber their parental taxa (Bansal et al., 2019; Geddes et al., 2021; Tangen et al., 2022)— suggesting that this prospective barrier is ineffective.

In this study, only two hybrids were observed in Europe, despite an extensive collecting trip that aimed to maximise the number of sampled hybrids (Ciotir et al., 2017). One of these hybrids was an F1 and had the *T. latifolia* plastome. Freeland et al., (2013) and Pieper et al., (2017) identified occasional *T. latifolia* cpDNA in hybrids. The presence of the *T. latifolia* plastome in some European hybrids raises the question of whether another mechanism of reproductive isolation between the two species in Europe could be related to the frequency of hybridisation events involving *T. latifolia* as maternal parent.

While *T. angustifolia* and *T. latifolia* are not sister species, they belong to two sister clades, and *T. latifolia* appears to have originated between 2 and 3 M generations ago (Aleman et al., 2025), i.e., ∼4 to 6 Mya, in line with Zhou et al (2018), assuming a 2- year generation time (Yeo, 1964). Our underestimation of their divergence time (∼1.25 M generations ago) is partially explained by the fact that they are not sister species; the missing branches between them might have inflated the species coalescence rates, making their divergence appear younger. Another factor contributing to this underestimate is that coalescent rates can anchor to periods of elevated demographic sizes (Wang et al., 2016); consistent with this, our divergence time estimation overlaps with a historical demographic peak observed in *T. latifolia* (Aleman et al., 2025). The successful posteriors goodness-of-fit (>0.05), similitude among the time of divergence and secondary contact estimations across all analyses and in this study (∼1.25 M generations ago), and the estimated time of secondary contact between the two species (∼400,000 generations ago)—coincident with the most recent demographic expansion experienced by *T. latifolia* (Aleman et al., 2025)—support the validity of our inferences.

Our estimated time of secondary contact between *T. angustifolia* and *T. latifolia* in North America was ∼430,000 generations ago, which predates the potential introduction of *T. angustifolia* to North America (Ciotir et al., 2013). When *T. angustifolia* entered North America, it would have carried the genomic signatures of contact with *T. latifolia* in Europe, which is how this date should be interpreted. It is expected that a coalescence approach would identify this historical contact in Europe, which reflects historical gene flow into *T. angustifolia* and is in line with Aleman et al. (2025), who observed introgressive hybridisation from *T. latifolia* into *T. angustifolia* in Europe.

*Typha latifolia* underwent a cycle of demographic declines and expansions between ∼2 M and 400,000 generations ago (equivalent to ∼4 M and 800,000 years) in North America and Europe (Aleman et al., 2025). Consistent with this, we estimated that *T. angustifolia* and *T. latifolia* came into contact ∼400,000 generations ago (equivalent to ∼800,000 years ago), suggesting the two species have been sympatric in Europe since at least the Middle Pleistocene, after the last expansion of *T. latifolia*. After coming into secondary contact, they likely hybridised and later developed reproductive barriers (Aleman et al., 2025).

A total of 47 candidate barrier loci between *T. angustifolia* and *T. latifolia* were identified in Europe, whereas none were identified in North America. This finding supports the hypothesis that reproductive barriers between the two species are stronger in Europe than in North America. A possible explanation for the rise of reproductive barriers between *T. angustifolia* and *T. latifolia* in Europe could be selection against hybrids (Hopkins, 2013; Pfennig, 2016)—future evidence will be necessary to support this idea. Identifying barrier loci in Europe but not in North America also supports the hypothesis that barriers to hybridisation could be stronger between species that have experienced prolonged periods in sympatry than those whose time in sympatry is short; further research could help determine if this is a general pattern of hybridisation. In flycatchers (Ficedula *albicollis* × *F. hypoleuca*) and oaks (*Quercus mongolica* × *Q. liaotungensis*), hybridisation rates are lower in older than younger areas of sympatry (Haavie et al., 2004; Liao et al., 2019). Similar patterns of reduced hybridisation in regions of prolonged sympatry have been observed in *Bombus* bumblebees and *Heliconius* butterflies, where selection strengthened reproductive barriers over time in contact (Christmas et al., 2021; Lewis et al., 2020).

Of the 47 barrier loci identified in Europe, 12 belonged to 10 genes associated with ubiquitin-mediated regulation and degradation (At4g11680) (Stone et al., 2005), protein modification and assembly (CASP1, CCB2, UGT80A2) (DeBolt et al., 2009; Lyska et al., 2007; Roppolo et al., 2011), membrane traffic and localization (VAMP714, ALMT12) (Gu et al., 2021; Sanderfoot, 2007; Sasaki et al., 2010), and important regulatory roles (IAA19, CINV1, PAIR2, and PDCD2). Notably, IAA19—which spanned three barrier loci (15 kb) and is under purifying selection between species in Europe—is involved in mediating auxin’s effects on plant growth (Kohno et al., 2012); CINV1 is involved in primary root elongation, lateral root formation, floral transition, and pollen development (Barratt et al., 2009); PAIR2 is essential for meiotic chromosome pairing (Nonomura et al., 2006); and PDCD2 is involved in apoptosis (Reape & McCabe, 2008). These genes are good candidates for causing reproductive barriers, supporting the role of barrier loci in contributing to reproductive isolation in Europe by impeding interbreeding events or making *T.* × *glauca* inviable or unfit.

The absence of barrier loci between *T. angustifolia* and *T. latifolia* in North America is consistent with the broad ranges of *T.* × *glauca* hybrids across the Great Lakes, Prairie Pothole, and Midwestern regions (reviewed in Bansal et al. 2019). On this continent, *T. latifolia* was potentially isolated since its arrival, between 5.7 and 3.5 Ma (Zhou et al., 2018), and until the potential arrival of *T. angustifolia* (Ciotir et al., 2013; Ciotir & Freeland, 2016). This time in isolation appears to have reduced the likelihood of reproductive barriers evolving between the two species, in contrast to Europe, where they have been sympatric for a prolonged time and reproductive barriers emerged after they hybridised in the past (Aleman et al., 2025). Alternatively, if barriers to hybridisation between the two species arose during a potential contact in Eurasia before *T. latifolia* arrived in North America, these barriers might have been lost during the time *T. latifolia* was isolated. In allopatry, selective pressures against (potentially maladaptive) hybrids are eliminated; without these pressures, local genetic differences driven by drift and adaptations might erode reproductive barriers that once prevented hybridisation (Coughlan & Matute, 2020). This dynamic—where reproductive isolation strengthens in sympatry due to selection against hybrids but weakens in allopatry due to its absence—is a common feature of species with cycles of isolation and contact (Kulmuni et al., 2020).

## Conclusions

Contributing to the results from Ciotir et al. (2013), Zhou et al. (2018), and Aleman et al. (2025), we propose that after diverging in allopatry due to drift-driven events ∼4 to 6 Mya, *T. angustifolia* and *T. latifolia* experienced secondary contact in Europe ∼400,000 generations ago (equivalent to ∼800,000 years ago), hybridised, and later developed reproductive isolation (reflected in 47 barrier loci). *Typha latifolia* arrived in North America from East Eurasia between 5.7 and 3.5 Ma and remained isolated until *T. angustifolia* arrived on this continent, potentially due to human dispersal. No barrier loci have arisen between the two species in North America. Future research could apply similar methods to *T. angustifolia* and *T. latifolia* in Eastern Asia—where they do not hybridise, and we predict more barrier loci should be found. Additionally, future research could investigate if selection against hybrids is the cause of reproductive barriers arising between *T. angustifolia* and *T. latifolia* in Europe. The lack of reproductive isolation between these two species in North America suggests that preventing future hybrid invasions will require limiting the movement of *Typha* and other allopatric species, which most likely lack reproductive barriers.

## Acknowledgements

We acknowledge that the laboratory procedures and data analyses for this study were conducted at Trent University, which is situated on the traditional territory of the Mississauga Anishinaabeg, to whom we extend our respect. We thank Polina Volkova and Tulsi Patel for their invaluable contributions to the laboratory and the field. This work was financially supported by the Natural Sciences and Engineering Research Council of Canada, and Alberto Aleman is funded by the Environmental and Life Sciences Graduate Program at Trent University. SHARCNET and Compute Canada provided computational resources for this study. Finally, we thank Erin Matula for her work on Figure 1.

## Author contribution statement

Conceptualisation, methodology, data analysis, writing, preparation of figures and tables: Aaron Shafer, Alberto Aleman, Joanna Freeland, and Marcel Dorken.

## Data availability

Data and scripts for this study can be found at https://gitlab.com/WiDGeT_TrentU/graduate_theses/-/tree/master/aleman/hdy and https://github.com/al-aleman/totoras_hdy.

