## Supplementary Tables S1 to S4 for "Time in sympatry correlates with the strength of reproductive isolation in hybridising *Typha*"

**Supplementary Table S1.** Identifiers, species memberships, approximate geographic coordinates, and continent of the samples collected in this study.

| Sampling ID | Species | Latitude | Longitude | Continent |
| --- | --- | --- | --- | --- |
| B12 | *T. × glauca* | 45.08 | -64.49 | North America |
| BRT2B | *T. angustifolia* | 44.74 | -63.24 | North America |
| BU14 | *T. × glauca* | 42.35 | -3.69 | Europe |
| C05TL | *T. latifolia* | 52.72 | -1.37 | Europe |
| CA13_TL | *T. latifolia* | 44.85 | 7.71 | Europe |
| CA3TL | *T. latifolia* | 47.68 | 22.46 | Europe |
| CC1_TL | *T. latifolia* | 50.77 | 2.31 | Europe |
| CC2_TL | *T. latifolia* | 50.77 | 2.31 | Europe |
| CIUU13TA | *T. angustifolia* | 43.76 | 24.93 | Europe |
| CL10TA | *T. angustifolia* | 50.77 | 2.31 | Europe |
| COTL6 | *T. latifolia* | 52.72 | -1.37 | Europe |
| CRR6 | *T. angustifolia* | 47.68 | 22.46 | Europe |
| CY1ATL | *T. latifolia* | 52.16 | 4.50 | Europe |
| CY6ATA | *T. angustifolia* | 51.43 | -0.11 | Europe |
| CY7ATA | *T. angustifolia* | 51.43 | -0.11 | Europe |
| CY7BTA | *T. angustifolia* | 51.43 | -0.11 | Europe |
| DAB16TA | *T. angustifolia* | 48.20 | 26.59 | Europe |
| E74B | *T. angustifolia* | 45.03 | -63.50 | North America |
| E91C | *T. angustifolia* | 45.03 | -63.50 | North America |
| EDD_44_1 | *T. latifolia* | 47.37 | -68.33 | North America |
| EDD_67_1 | *T. latifolia* | 47.37 | -68.33 | North America |
| EDW_2_15 | *T. latifolia* | 47.37 | -68.33 | North America |
| EDW_3_6 | *T. latifolia* | 47.37 | -68.33 | North America |
| EL14_TA | *T. latifolia* | 52.44 | 5.85 | Europe |
| ELTA_01 | *T. angustifolia* | 52.44 | 5.85 | Europe |
| ELTA_08 | *T. angustifolia* | 52.44 | 5.85 | Europe |
| FRD47_2 | *T. latifolia* | 45.96 | -66.64 | North America |
| GORE_3 | *T. latifolia* | 43.68 | -70.44 | North America |
| G_II3 | *T. latifolia* | 49.11 | 11.93 | Europe |
| HAD_3_1 | *T. angustifolia* | 44.60 | -63.55 | North America |
| HAW_03_12 | *T. latifolia* | 44.60 | -63.55 | North America |
| HAW_10_9 | *T. latifolia* | 44.60 | -63.55 | North America |
| HM3TL | *T. latifolia* | 47.51 | 19.04 | Europe |
| HOR_6 | *T. latifolia* | 48.14 | 26.51 | Europe |
| HUF_13TA | *T. angustifolia* | 47.69 | 17.65 | Europe |
| HUG_13TA | *T. angustifolia* | 47.69 | 17.65 | Europe |
| HUS_10TL | *T. latifolia* | 47.51 | 19.04 | Europe |
| HUS_TA1 | *T. angustifolia* | 47.51 | 19.04 | Europe |
| HUS_TL1 | *T. latifolia* | 47.51 | 19.04 | Europe |
| IC17TA | *T. angustifolia* | 43.57 | 4.32 | Europe |
| ICTD5 | *T. angustifolia* | 43.57 | 4.32 | Europe |
| IP2 | *T. latifolia* | 44.46 | -64.32 | North America |
| IR_01 | *T. × glauca* | 44.97 | -64.06 | North America |
| IR_105 | *T. × glauca* | 44.97 | -64.06 | North America |
| IR_11 | *T. angustifolia* | 44.97 | -64.06 | North America |
| IR_119 | *T. × glauca* | 44.97 | -64.06 | North America |
| IR_25 | *T. latifolia* | 44.97 | -64.06 | North America |
| IR_36 | *T. latifolia* | 44.97 | -64.06 | North America |
| IR_39 | *T. latifolia* | 44.97 | -64.06 | North America |
| IR_41 | *T. latifolia* | 44.97 | -64.06 | North America |
| KL14_TL | *T. latifolia* | 46.64 | 14.31 | Europe |
| KL1TL | *T. latifolia* | 46.64 | 14.31 | Europe |
| KL28_TL | *T. latifolia* | 46.64 | 14.31 | Europe |
| KL_TH1 | *T. latifolia* | 46.64 | 14.31 | Europe |
| LiETL_11 | *T. latifolia* | 42.21 | 2.61 | Europe |
| M114 | *T. × glauca* | 45.08 | -64.49 | North America |
| M23 | *T. latifolia* | 45.08 | -64.49 | North America |
| M30 | *T. latifolia* | 45.08 | -64.49 | North America |
| M37 | *T. latifolia* | 45.08 | -64.49 | North America |
| M53 | *T. latifolia* | 45.08 | -64.49 | North America |
| M85 | *T. latifolia* | 45.08 | -64.49 | North America |
| MOD_01_0 | *T. latifolia* | 46.98 | -70.55 | North America |
| MOD_02_1 | *T. latifolia* | 46.98 | -70.55 | North America |
| MOD_04_1 | *T. latifolia* | 46.98 | -70.55 | North America |
| MOD_05_2 | *T. latifolia* | 46.98 | -70.55 | North America |
| MOD_07_0 | *T. angustifolia* | 46.98 | -70.55 | North America |
| MOD_08_1 | *T. latifolia* | 46.98 | -70.55 | North America |
| MOD_09_0 | *T. angustifolia* | 46.98 | -70.55 | North America |
| MOD_14_0 | *T. latifolia* | 46.98 | -70.55 | North America |
| MOD_14_3 | *T. angustifolia* | 46.98 | -70.55 | North America |
| MOD_15_1 | *T. latifolia* | 46.98 | -70.55 | North America |
| MOD_16_1 | *T. latifolia* | 46.98 | -70.55 | North America |
| MOD_17_1 | *T. latifolia* | 46.98 | -70.55 | North America |
| MOD_18_0 | *T. latifolia* | 46.98 | -70.55 | North America |
| MOD_25_2 | *T. latifolia* | 46.98 | -70.55 | North America |
| MOD_26_2 | *T. latifolia* | 46.98 | -70.55 | North America |
| MOD_29_1 | *T. latifolia* | 46.98 | -70.55 | North America |
| MOD_33_0 | *T. × glauca* | 46.98 | -70.55 | North America |
| MOD_35_3 | *T. latifolia* | 46.98 | -70.55 | North America |
| MOD_37_1 | *T. × glauca* | 46.98 | -70.55 | North America |
| MOD_38_1 | *T. × glauca* | 46.98 | -70.55 | North America |
| MOD_39_0 | *T. × glauca* | 46.98 | -70.55 | North America |
| MOD_40_0 | *T. × glauca* | 46.98 | -70.55 | North America |
| MOD_42_3 | *T. × glauca* | 46.98 | -70.55 | North America |
| MOD_45_2 | *T. latifolia* | 46.98 | -70.55 | North America |
| MOD_47_1 | *T. angustifolia* | 46.98 | -70.55 | North America |
| MOD_50_1 | *T. × glauca* | 46.98 | -70.55 | North America |
| MOD_52_0 | *T. × glauca* | 46.98 | -70.55 | North America |
| MOD_54_0 | *T. latifolia* | 46.98 | -70.55 | North America |
| MOD_55_0 | *T. × glauca* | 46.98 | -70.55 | North America |
| MOD_56_1 | *T. angustifolia* | 46.98 | -70.55 | North America |
| MOD_57_0 | *T. latifolia* | 46.98 | -70.55 | North America |
| MOD_58_0 | *T. × glauca* | 46.98 | -70.55 | North America |
| MOD_59_1 | *T. × glauca* | 46.98 | -70.55 | North America |
| MOD_60_0 | *T. × glauca* | 46.98 | -70.55 | North America |
| MOD_61_0 | *T. × glauca* | 46.98 | -70.55 | North America |
| MOD_62_1 | *T. × glauca* | 46.98 | -70.55 | North America |
| MOD_63_0 | *T. × glauca* | 46.98 | -70.55 | North America |
| ORP_TI | *T. latifolia* | 43.28 | -2.13 | Europe |
| OR_TI2 | *T. latifolia* | 43.28 | -2.13 | Europe |
| OR_TL5 | *T. latifolia* | 43.28 | -2.13 | Europe |
| P4 | *T. latifolia* | 45.41 | -64.33 | North America |
| P6 | *T. latifolia* | 45.41 | -64.33 | North America |
| P9 | *T. latifolia* | 45.41 | -64.33 | North America |
| PID_04_3 | *T. × glauca* | 43.84 | -79.10 | North America |
| PID_08_5 | *T. × glauca* | 43.84 | -79.10 | North America |
| PID_09_0 | *T. × glauca* | 43.84 | -79.10 | North America |
| PID_10_1 | *T. × glauca* | 43.84 | -79.10 | North America |
| PIW_01_0 | *T. angustifolia* | 43.84 | -79.10 | North America |
| PIW_04_6 | *T. angustifolia* | 43.84 | -79.10 | North America |
| PIW_06_6 | *T. angustifolia* | 43.84 | -79.10 | North America |
| PIW_07_1 | *T. angustifolia* | 43.84 | -79.10 | North America |
| PIW_09_1 | *T. angustifolia* | 43.84 | -79.10 | North America |
| PIW_10_2 | *T. angustifolia* | 43.84 | -79.10 | North America |
| PIW_12_3 | *T. × glauca* | 43.84 | -79.10 | North America |
| PIW_13_0 | *T. angustifolia* | 43.84 | -79.10 | North America |
| PIW_14_3 | *T. angustifolia* | 43.84 | -79.10 | North America |
| PIW_15_3 | *T. angustifolia* | 43.84 | -79.10 | North America |
| PIW_16_2 | *T. angustifolia* | 43.84 | -79.10 | North America |
| PIW_20_6 | *T. angustifolia* | 43.84 | -79.10 | North America |
| PIW_21_3 | *T. angustifolia* | 43.84 | -79.10 | North America |
| PIW_22_6 | *T. angustifolia* | 43.84 | -79.10 | North America |
| PIW_23_3 | *T. angustifolia* | 43.84 | -79.10 | North America |
| PIW_24_3 | *T. angustifolia* | 43.84 | -79.10 | North America |
| PIW_25_3 | *T. angustifolia* | 43.84 | -79.10 | North America |
| PWE19TL | *T. latifolia* | 51.62 | -3.94 | Europe |
| RGT2 | *T. angustifolia* | 45.48 | -74.30 | North America |
| SADT_03 | *T. angustifolia* | 45.90 | -64.39 | North America |
| SAD_10_3 | *T. latifolia* | 45.90 | -64.39 | North America |
| SAD_17_3 | *T. latifolia* | 45.90 | -64.39 | North America |
| SAD_25_0 | *T. latifolia* | 45.90 | -64.39 | North America |
| SCH9TA | *T. latifolia* | 47.65 | 26.22 | Europe |
| SP2_TA | *T. angustifolia* | 52.41 | 4.68 | Europe |
| SR2_TA | *T. × glauca* | 52.41 | 4.68 | Europe |
| TA_01 | *T. angustifolia* | 51.48 | 0.61 | Europe |
| TB7 | *T. latifolia* | 44.39 | -64.25 | North America |
| TMO2_7 | *T. latifolia* | 46.09 | -64.78 | North America |
| Vikram_202 | *T. × glauca* | 44.30 | -78.32 | North America |
| Vikram_237 | *T. × glauca* | 44.30 | -78.32 | North America |
| Vikram_25 | *T. angustifolia* | 44.30 | -78.32 | North America |
| Vikram_28 | *T. × glauca* | 44.30 | -78.32 | North America |
| Vikram_33 | *T. angustifolia* | 44.30 | -78.32 | North America |
| Vikram_73 | *T. angustifolia* | 44.30 | -78.32 | North America |
| Vikram_76 | *T. latifolia* | 44.30 | -78.32 | North America |
| Vikram_86 | *T. latifolia* | 44.30 | -78.32 | North America |
| Vikram_94 | *T. angustifolia* | 44.30 | -78.32 | North America |
| Vikram_95 | *T. latifolia* | 44.30 | -78.32 | North America |
| WBD20_1 | *T. angustifolia* | 44.28 | -84.23 | North America |
| WBD_22_5 | *T. × glauca* | 44.28 | -84.23 | North America |
| WBD_27_4 | *T. angustifolia* | 44.28 | -84.23 | North America |
| WBD_28_3 | *T. angustifolia* | 44.28 | -84.23 | North America |
| WBD_29_2 | *T. × glauca* | 44.28 | -84.23 | North America |
| WBD_32_1 | *T. angustifolia* | 44.28 | -84.23 | North America |
| WBD_33_1 | *T. angustifolia* | 44.28 | -84.23 | North America |
| WBD_34_1 | *T. × glauca* | 44.28 | -84.23 | North America |
| WBD_36_2 | *T. × glauca* | 44.28 | -84.23 | North America |
| WBW5_4 | *T. latifolia* | 44.28 | -84.23 | North America |
| WBW_14_4 | *T. × glauca* | 44.28 | -84.23 | North America |
| WBW_16_2 | *T. × glauca* | 44.28 | -84.23 | North America |
| WBW_18_6 | *T. × glauca* | 44.28 | -84.23 | North America |

**Supplementary Table S2.** Prior bounds of the divergence times and effective population sizes used during the reconstruction of the demographic and divergence histories of *T. angustifolia* and *T. latifolia* in North America and Europe.

|  | North America | Europe | *T. angustifolia* (North America) –  *T. latifolia* (Europe) | *T. angustifolia* (Europe) –  *T. latifolia*  (North America) |
| --- | --- | --- | --- | --- |
| Minimum effective size (N_e_) | 1,071 | 1,429 | 1,429 | 1,071 |
| Minimum effective size (N_e_) | 608,929 | 652,036 | 608,929 | 652,036 |
| Maximum divergence time (in generations) | 864,285 | 801,482 | 834,821 | 830,946 |
| Maximum divergence time (in generations) | 1,728,571 | 1,602,965 | 1,669,642 | 1,661,892 |

**Table S3.** Likelihood of the different scenarios tested during the reconstruction of the demographic history of *T. angustifolia* and *T. latifolia* in North America and Europe. Scenarios with the highest likelihood are denoted in **bold**. AM = Ancestral migration; IM = Isolation–Migration; SC = Secondary contact; SI = Strict Isolation; 1M = Rates of interspecific gene flow remained constant over time; 2M = Rates of interspecific gene flow changed over time; 1N = Effective sizes (N_e_) remained constant over time; 2N = Effective sizes (N_e_) changed over time.

|  | North America | Europe | *T. angustifolia*  (North America) –  *T. latifolia* (Europe) | *T. angustifolia* (Europe) –  *T. latifolia*  (North America) |
| --- | --- | --- | --- | --- |
| AM_1M_1N | 0.00131801 | 0.00275322 | 0.00158906 | 0.0007579 |
| AM_1M_2N | 0.01451122 | 0.01326868 | 0.00758491 | 0.01287907 |
| AM_2M_1N | 0.00151503 | 0.0095897 | 0.01217523 | 0.00171429 |
| AM_2M_2N | 0.01475842 | 0.01662897 | 0.00881292 | 0.01834401 |
| IM_1M_1N | 0.04620111 | 0.04382832 | 0.0344655 | 0.0420938 |
| IM_1M_2N | 0.12083723 | 0.10250842 | 0.07366645 | 0.09782658 |
| IM_2M_1N | 0.08616011 | 0.09540489 | 0.08519177 | 0.10659093 |
| IM_2M_2N | **0.21147222** | 0.16850417 | 0.1730375 | **0.18919861** |
| SC_1M_1N | 0.08448302 | 0.09026685 | 0.10169659 | 0.07249265 |
| SC_1M_2N | 0.16397194 | 0.15623251 | 0.1673672 | 0.18099716 |
| SC_2M_1N | 0.08190043 | 0.10936329 | 0.09894899 | 0.10191291 |
| SC_2M_2N | 0.16227781 | **0.18477438** | **0.23111132** | 0.16810195 |
| SI_1N | 0.0003016 | 0.00097208 | 0.00126732 | 0.00030913 |
| SI_2N | 0.01029183 | 0.00590454 | 0.00308525 | 0.00678103 |

**Table 2.** Characterisation of barrier loci between *T. angustifolia* and *T. latifolia* in Europe and between species from different continents. No barrier loci were identified in North America. Unless otherwise indicated in the observations, barrier loci were found in Europe. BLAST = Genes predicted by BLAST (Camacho et al. 2009).

| Chromosome | Coordinates | Annotation/Predicted protein | Reference | Observations |
| --- | --- | --- | --- | --- |
| 1 | 6585000 – 6590000 | At4g11680 | (Stone et al. 2005) | – |
| 1 | 12400000 – 12405000 | Cinv1 | (Barratt et al. 2009) | Positive selection in *T. latifolia* |
| 1 | 13925000 – 13930000 | AP2-like ethylene-responsive transcription factor AIL5 | BLAST | F_ST_ outlier |
| 1 | 22255000 – 22260000 | Uncharacterized protein | BLAST | F_ST_ outlier |
| 1 | 32070000 – 32075000 | Serine/threonine-protein kinase D6PK-like | BLAST | – |
| 2 | 65000 – 70000 | Dof zinc finger protein DOF2.1-like | BLAST | F_ST_ outlier |
| 2 | 13045000 – 13050000 | Ccb2 | (Lyska et al. 2007) | F_ST_ outlier |
| 2 | 16240000 – 16245000 | Ugt80a2 | (DeBolt et al. 2009) | F_ST_ outlier |
| 2 | 17600000 – 17605000 | Uncharacterized protein | BLAST | – |
| 2 | 17685000 – 17690000 | Uncharacterized protein | BLAST | – |
| 2 | 21520000 – 21525000 | Uncharacterized protein | BLAST | – |
| 2 | 25195000 – 25200000 | Casp1 | (Roppolo et al. 2011) | – |
| 3 | 650000 – 655000 | CBBY-like protein | BLAST | F_ST_ outlier |
| 3 | 8840000 – 8845000 | Glutathione gamma-glutamylcysteinyltransferase 1-like | BLAST | – |
| 3 | 20660000 – 20665000 | Transcription factor KUA1-like | BLAST | – |
| 3 | 22825000 – 22830000 | Uncharacterized protein | BLAST | – |
| 3 | 23070000 – 23075000 | Xyloglucan endotransglucosylase protein 6-like | BLAST | – |
| 4 | 1400000 – 1405000 | F-box/LRR-repeat protein At3g26922-like | BLAST | – |
| 4 | 10560000 – 10565000 | Polyol transporter 5-like | BLAST | – |
| 4 | 23330000 – 23335000 | UPF0481 protein At3g47200 – like | BLAST | F_ST_ and d_XY_ outlier |
| 5 | 8975000 – 8980000 | Uncharacterized protein | BLAST | – |
| 6 | 820000 – 825000 | Uncharacterized protein | BLAST | – |
| 7 | 3645000 – 3650000 | – | – | F_ST_ outlier |
| 7 | 12275000 – 12280000 | Probable disease resistance protein At4g14610 | BLAST | – |
| 7 | 12320000 – 12325000 | Uncharacterized protein | BLAST | – |
| 8 | 860000 – 865000 | Uncharacterized protein | BLAST | – |
| 8 | 3230000 – 3235000 | Uncharacterized protein | BLAST | – |
| 9 | 11845000 – 11850000 | Pair2 | (Nonomura et al. 2006) | – |
| 9 | 13585000 – 13590000 | Uncharacterized protein | BLAST | – |
| 10 | 6130000 – 6135000 | – | – | – |
| 11 | 725000 – 730000 | SWI/SNF complex component SNF12 homolog | BLAST | – |
| 11 | 1975000 – 1980000 | Uncharacterized lncRNA | BLAST | – |
| 11 | 4780000 – 4785000 | Uncharacterized lncRNA | BLAST | Only in *T. angustifolia* (North America) vs. *T. latifolia* (Europe) |
| 11 | 5550000 – 5555000 | – | – | Both in (i) Europe and (ii)  *T. angustifolia* (Europe) vs.  *T. latifolia* (North America) |
| 11 | 5880000 – 5885000 | Uncharacterized lncRNA | BLAST | Both in (i) Europe and (ii)  *T. angustifolia* (North America) vs.  *T. latifolia* (Europe) |
| 11 | 10935000 – 10940000 | Disease resistance protein RGA2-like | BLAST | – |
| 12 | 7595000 – 7600000 | Tubulin beta-3 chain | BLAST | – |
| 13 | 4645000 – 4650000 | Gamma-tubulin complex component 2 LAT | BLAST | – |
| 13 | 9235000 – 9240000 | Pdcd2 | (Baron et al. 2010) | – |
| 13 | 11910000 – 11925000 | Iaa19 | (Liscum and Reed 2002) | Purifying selection between species |
| 13 | 12670000 – 12675000 | Almt12 | (Sasaki et al. 2010) | – |
| 14 | 70000 – 75000 | ABC transporter C family member 5-like | BLAST | – |
| 14 | 5990000 – 5995000 | Receptor-like protein EIX2 | BLAST | – |
| 14 | 12090000 – 12095000 | Uncharacterized protein (LOC140763314), mRNA | BLAST | Only in *T. angustifolia* (North America) vs. *T. latifolia* (Europe) |
| 15 | 9910000 – 9915000 | Uncharacterized lncRNA | BLAST | – |
| 15 | 10745000 – 10750000 | Phosphatidylinositol/phosphatidylcholine transfer protein SFH9-like | BLAST | – |
| 15 | 11705000 – 11710000 | Probable E3 ubiquitin-protein ligase ARI8 | BLAST | F_ST_ outlier |
